# Long-term warming effects on the microbiome and nitrogen fixation of a common moss species in sub-Arctic tundra

**DOI:** 10.1101/838581

**Authors:** Ingeborg J. Klarenberg, Christoph Keuschnig, Ana J. Russi Colmenares, Denis Warshan, Anne D. Jungblut, Ingibjörg S. Jónsdóttir, Oddur Vilhelmsson

## Abstract

1. Bacterial communities form the basis of biogeochemical processes and determine plant growth and health. Mosses, an abundant plant group in Arctic ecosystems, harbour diverse bacterial communities that are involved in nitrogen fixation and carbon cycling. Global climate change is causing changes in aboveground plant biomass and shifting species composition in the Arctic, but little is known about the response of moss microbiomes.
2. Here, we studied the total and potentially active bacterial community associated with *Racomitrium lanuginosum*, in response to 20-year *in situ* warming in an Icelandic heathland. We evaluated the effect of warming and warming-induced shrub expansion on the moss bacterial community composition and diversity, *nifH* gene abundance and nitrogen-fixation rates.
3. Warming changed both the total and the potentially active bacterial community structure, while litter abundance only affected the total bacterial community structure. The relative abundance of Proteobacteria increased, while the relative abundance of Cyanobacteria and Acidobacteria decreased. *NifH* gene abundance and nitrogen-fixation rates were negatively affected by litter and *Betula nana* abundance, respectively. We also found shifts in the potentially nitrogen-fixing community, with *Nostoc* decreasing and non-cyanobacterial diazotrophs increasing in relative abundance. Our data suggests that the moss microbial community including the potentially nitrogen-fixing taxa is sensitive to future warming.
4. *Synthesis.* Long-term warming led to a shift in moss-associated bacterial community composition, while the abundance of nitrogen-fixing bacteria and nitrogen-fixation rates were negatively affected by increased litter and *Betula nana* abundance respectively. Warming and increased shrub abundance as a result of warming can affect moss-associated bacterial communities and nitrogen fixation rates in tundra ecosystems.

## Introduction

Temperature in high-latitude regions is rising twice as fast as elsewhere (IPCC, 2019), which is predicted to have large impacts on Arctic ecosystems, for instance by altering species distributions and interactions (Van der Putten, 2012; Wookey et al., 2009). One such interaction that might be affected by warming is the association between mosses and bacterial communities as well as related ecosystem processes such as pedogenesis, carbon (C) cycling, and nitrogen (N) cycling.

Bryophytes, mosses in particular, comprise a large component of the vegetation in many high-latitude ecosystems (Longton, 1992). They play important roles in biogeochemical cycles by forming a C sink via their slow decomposition rates, by accounting for up to 7% of terrestrial net primary productivity and by supporting up to half of the terrestrial N_2_-fixation (Cornelissen et al., 2007; 2012; Porada et al., 2013; Turetsky, 2003; Turetsky et al., 2012). Most mosses consist of a upper living segment with photosynthetic tissue and a lower decaying dead segment and thus link above-ground and belowground processes (Whiteley & Gonzalez, 2016). Mosses provide a habitat for a range of microbiota, microfauna and mesofauna (Lindo & Gonzalez, 2010). These moss-associated microorganisms are involved in the decomposition of dead moss tissue (Kulichevskaya et al., 2007) and some of them are active diazotrophs (Chen et al., 2019). N_2_-fixation by moss-associated Cyanobacteria, the best studied of these diazotrophs, was shown to directly increase moss growth rates (Berg et al., 2013) and thereby control C sequestration in moss tissues. Moss-associated diazotrophy is also an important source of new available N in boreal and Arctic ecosystems (DeLuca et al., 2002; Rousk et al., 2017). In order to understand the implications of climate change for the role of mosses in ecosystem C and N cycling, we need to understand how moss-associated microbial communities react to elevated temperatures.

The bacterial community composition of mosses is species specific and influenced by environmental factors such as pH and nutrient availability (Bragina, Berg, et al., 2012; Holland Moritz et al., 2018; Tang et al., 2016). While Cyanobacteria have received most of the attention for their N_2_-fixing capability (Berg et al., 2013; Gentili et al., 2005; Ininbergs et al., 2011; Lindo et al., 2013; Rousk et al., 2013; Stewart, Lamb, et al., 2011; Warshan et al., 2016, 2017), mosses harbour diverse bacterial communities. Commonly found phyla associated with mosses include Acidobacteria, Actinobacteria, Armatimonadetes, Bacteroidetes, Cyanobacteria, Planctomycetes, Proteobacteria and Verrucomicrobia (Kostka et al., 2016; Tang et al., 2016), and their potential functions include N_2_-fixation (Bragina, Maier, et al., 2012), anoxygenic phototrophy (Holland Moritz et al., 2018) and freeze protection (Raymond, 2016). The bacterial community composition of mosses has primarily been studied for peat and feather mosses, but we know little about the bacterial communities of other moss species. For instance, little is known about the bacterial community associated with ecologically important moss species such as *Racomitrium lanuginosum* (Hedw.) Brid. This moss species has a wide distribution at high altitudes in temperate regions of the Northern and Southern Hemisphere and at low altitudes in the Arctic (Jonsdottir et al., 1995; Tallis, 1995). It is a dominant species in many Icelandic ecosystems, forming dense mats where conditions are favourable for colonisation and growth (Bjarnason, 1991; Ingimundardóttir et al., 2014; Tallis, 1958).

Despite the importance of microbial communities for plant functioning and ecosystem processes, the long-term effect of warming on moss microbial communities has received little attention. Two studies describing the effect of four weeks to two years warming-related changes in peat moss bacterial community composition, reported a decrease in overall bacterial and diazotrophic diversity with higher temperatures *in situ* and under laboratory conditions (Carrell et al., 2019; Kolton et al., 2019). Whether this warming-induced decrease in diversity also holds for bacterial communities associated with other moss species in high latitudes is unknown. Moreover, decades-long-warming effects on moss-associated bacterial communities have yet to be explored.

Nonetheless, the effect of warming on some high-latitude plant communities has been better documented, where for instance ambient and experimental warming (ranging from 5-43 years) in tundra heaths have resulted in shrub expansion (Bjorkman et al., 2020; Myers-Smith et al., 2011; Myers[Smith et al., 2019). The increase in deciduous dwarf shrubs, for example *Betula nana*, led to an increase in the quantity of relatively high quality litter, resulting in a faster turnover of the overall leaf litter C and N (McLaren et al., 2017). This warming-induced change in litter quality and nutrient cycling might also affect the composition of microbial communities (Deslippe et al., 2012). Indeed, changing litter inputs can consequently lead to shifts in moss microbiomes (Jean et al., 2020). The increase in labile shrub litter may lead to an increase in copiotrophic taxa and decrease in oligotrophic taxa (Fierer et al., 2007; Wallenstein et al., 2007). Warming might thus also, indirectly, via a change in leaf litter quality and quantity resulting from increasing shrub biomass, lead to changes in the bacterial communities associated with the moss layer. Changes in bacterial community composition could consequently affect N_2_-fixation rates (Wu et al., 2020). In addition, N_2_-fixation rates can be expected to increase with temperature, as metabolic process rate in microorganisms increases with temperature and the enzyme nitrogenase is more active at higher temperatures than average Arctic temperatures (Houlton et al., 2008). Temperature-induced drought, however, can inhibit N_2_-fixation rates, especially cyanobacterial N_2_-fixation (Rousk et al., 2014, 2015, 2018; Stewart, Coxson, et al., 2011; Stewart et al., 2014; Stewart, Lamb, et al., 2011; Whiteley & Gonzalez, 2016; Zielke et al., 2005). Indirect effects of temperature on N_2_-fixation rates might also be related to physiological adaptation of diazotrophic communities (Whiteley & Gonzalez, 2016), or shifts to a species composition better suited to the new conditions (Deslippe et al., 2005; Rousk et al., 2018; Rousk & Michelsen, 2017). Warming-induced changes in bacterial species composition could potentially feedback to the abundance, diversity and/or N_2_-fixation activity of diazotrophs, through alteration of biotic interactions between bacteria e.g. competition and/or cooperation (Ho et al., 2016). The increase in shrubs might also affect N_2_-fixation rates, either negatively via an increase in shading leading to an decrease in N_2_-fixation rates, or either inhibit or promote N_2_-fixation depending on the nutrient content of the litter (Rousk & Michelsen, 2017; Sorensen & Michelsen, 2011).

In this study we investigated how two decades of experimental warming with open top chambers impact the bacterial community and N_2_-fixation rates associated with the prevailing moss *R. lanuginosum* (Hedw.) Brid in a subarctic-alpine dwarf shrub heath in northern Iceland, dominated by *B. nana*.

We hypothesised that long-term warming directly and/or indirectly via the warming-induced increase in labile *B. nana* litter (1) leads to a shift in bacterial community composition with a decrease in bacterial diversity and (2) leads to a decrease in oligotrophic taxa and an increase in copiotrophic taxa. Further, we hypothesised (3) that changes in N_2_-fixation rates will depend on the combination of the direct effect of warming leading to an increase in N_2_-fixation rates and indirect effects of warming. These indirect effects include shading leading to a decrease in N_2_-fixation rates; increased litter leading to an increase or a decrease in N_2_-fixation rates; and/or changes in the bacterial community that could mediate the effects of warming, shading and/or litter on N_2_-fixation rates. To address these hypotheses, we sampled *R. lanuginosum* in a warming simulation experiment in the northwest highlands of Iceland that has been running for 20 years (Jonsdottir et al., 2005). We assessed the associated bacterial community structure by 16S rRNA gene and rRNA amplicon sequencing, N_2_-fixation rates with acetylene reduction assays (ARA), and N_2_-fixation potential using quantitative PCR (qPCR) of the *nifH* gene encoding the iron-protein component of the nitrogenase.

## Methods

### Field site and experimental design

The sampling was conducted in permanent plots of a long-term warming simulation experiment at Auðkúluheiði in the northwest highlands of Iceland (65°16’N, 20°15’W, 480 m above sea level). The site is a part of the International Tundra Experiment (ITEX; Henry and Molau 1997) and according to Köppen’s climate definitions, the sampling site is situated within the lower Arctic (Köppen, 1931). The vegetation has been characterized as a relatively species-rich dwarf shrub heath, with *B. nana* being the most dominant vascular species and *R. lanuginosum* and *Cetraria islandica* as the dominating moss and lichen species (Jonsdottir et al., 2005). The experimental site has been fenced off since 1996 to prevent sheep from disturbing the experiment.

Ten plot pairs of 75x75 cm were selected and one of the plots in each pair was randomly assigned to a warming treatment while the other served as a control. Open top plexiglass chambers (OTCs) were set up in August 1996 and 1997 to simulate a warmer summer climate and have been in place throughout the year ever since (Hollister & Webber, 2000; Jonsdottir et al., 2005). The temperature in the OTCs was on average 1.4 °C higher in June 2016 to August 2016 and 0.22 °C higher from August 2018 to June 2019 (Table S1). Relative humidity was -3 % lower in the OTCs in June 2016 to August 2016 (Table S1).

The vegetation responses were monitored by a detailed vegetation analysis after peak biomass at a few year intervals using the point intercept method following standard protocols of the International Tundra Experiment (Molau & Mølgaard, 1996): 100 points per plot, all hits (intercepts) per species recorded in each point through the canopy; relates to biomass) (Jonsdottir et al., 2005). In this study we use data from August 2014 on abundance (total number of hits per plot) for *R. lanuginosum*, *B. nana* and litter to test hypotheses 1-3 (Table S2). In 2014 the abundance of *R. lanuginosum* was on average 0.8 times lower in the warmed plots than control plots, but not significantly, while the abundance of *B. nana* was 2.5 times greater in the warm plots on average and litter was 2.7 times greater (Table S2).

### RNA and DNA extraction and sequencing

To assess overall bacterial community structure and bacterial diversity (hypothesis 1 and 2) associated with *R. lanuginosum* we collected moss shoots, extracted DNA and RNA and used 16S rRNA gene amplicon sequencing. For RNA and DNA extraction we collected *R. lanuginosum* moss shoots in June 2017. Per warmed (OTC) and control plot, five moss shoots were collected with sterile tweezers. In total 50 OTC and 50 control samples were collected. The moss shoots were immediately soaked in RNAlater (Ambion) to prevent RNA degradation and kept cool until storage at -80 °C. Prior to extraction, the samples were rinsed with RNase free water to remove soil particles and RNAlater and ground for six minutes using a Mini-Beadbeater and two sterile steel beads. RNA and DNA were extracted simultaneously using the RNeasy PowerSoil Total RNA Kit (Qiagen) and the RNeasy PowerSoil DNA Elution Kit (Qiagen), following the manufacturer’s instructions. DNA and RNA concentrations were determined with a Qubit Fluorometer (Life Technologies) and quality was assessed with a NanoDrop (NanoDrop Technologies) and Bioanalyzer (Agilent Technologies). cDNA was synthesized using the High-Capacity cDNA Reverse Transcription Kit (Thermofisher) following the manufacturer’s instructions and quantified on a Qubit Fluorometer (Life Technologies). All DNA extractions (100 samples) were used for qPCR. From all DNA and cDNA samples, we selected 48 DNA samples (24 from each treatment) and 48 cDNA samples (24 from each treatment) for sequencing based on RNA and DNA quality and quantity. Library preparation and sequencing of the V3-V4 region of the 16S rRNA gene on an Illumina MiSeq platform (2 x 300 bp) was performed by Macrogen, Seoul, using MiSeq v3 reagents and the primer pair 337F/805R and the PCR conditions described in (Klindworth et al., 2013).

### Sequence processing

In order to obtain high-resolution data and to better discriminate ecological patterns, we processed the raw sequences using the DADA2 (version 1.12.1) pipeline (Callahan et al., 2016, 2017), which does not cluster sequences into operational taxonomic units (OTUs), but uses exact sequences or amplicon sequence variants (ASVs). Forward reads were truncated at 260 bp and reverse reads at 250 bp. Assembled ASVs were assigned taxonomy using the Ribosomal Database Project (RDP) naïve Bayesian classifier (Q. Wang et al., 2007) in DADA2 and the SILVA_132 database (Quast et al., 2013). We removed samples with less than 10.000 non-chimeric sequences (11 samples) and we removed ASVs assigned to chloroplasts and mitochondria, singletons, as well as ASVs present in only one sample. In total, for 85 samples, 3598 ASVs remained with an average read size of 448 bp after DADA2. To account for uneven sequencing depths, the data were normalised using cumulative-sum scaling (CSS) (Paulson et al., 2013). The 16S rRNA gene based community is hereafter sometimes referred to as the ‘total bacterial community’ and the 16S rRNA (cDNA) based community is hereafter referred to as the ‘potentially metabolically active bacterial community’, acknowledging that 16S rRNA is not a direct indicator of activity but rather protein synthesis potential (Blazewicz et al., 2013). Raw sequences are available in the European Nucleotide Archive under accession number PRJEB40635.

### Quantitative real-time PCR of nifH and 16S rRNA genes

We used the DNA samples (100 samples (50 control and 50 OTC samples)) for quantification of *nifH* and 16S rRNA genes (to test hypothesis 3). This was performed by quantitative PCR (Corbett Rotor-Gene) using the primer set PolF/PolR and 341F/534R respectively (Poly et al., 2001). The specificity of the *nifH* primers for our samples was confirmed by SANGER sequencing of 10 clone fragments. Standards for *nifH* reactions were obtained by amplifying one cloned *nifH* sequence with flanking regions of the plasmid vector (TOPO TA cloning Kit, Invitrogen). Standard curves were obtained by serial dilutions (E = 0.9 – 1.1, R^2^ = > 0.99 for all reactions). Each reaction had a volume of 20 µL, containing 1x QuantiFast SYBR Green PCR Master Mix (Qiagen), 0.2 µL of each primer (10 µM), 0.8 µL BSA (5 µg/µL), 6.8 µL RNase free water and 2 µL template. The cycling program was 5 min at 95 °C, 30 cycles of 10 s at 95 °C and 30 s at 60 °C.

### Acetylene reduction assays

We used acetylene reduction assays (ARA) to estimate N_2_-fixation rate to test hypothesis 3. We followed the procedure described in DeLuca et al. (2002) and Zackrisson et al. (2004). We collected three moss shoots of 5 cm length per control plot and OTC in June and August 2014. The three shoots per plot were analysed separately. The moss shoots were placed in 20 mL vials with 2 mL deionized water. Moss shoots were acclimated in a growth mol μm^−2^ s^−1^ PAR. 10% of the headspace was replaced by acetylene. After an additional 24 h of incubation in the growth chamber under the same conditions, acetylene reduction and ethylene production were measured by gas chromatography.

### Statistical analysis

All statistical analyses were performed in R (version 3.6.3). Richness (number of ASVs) and Shannon diversity were calculated with the R packages ‘vegan’ (version 2.5-4) (Oksanen et al., 2013) and ‘phyloseq’ (version 1.28.0) (McMurdie & Holmes, 2013). Differences in N_2_-fixation rates, 16S rRNA and *nifH* gene abundance, ASV richness and Shannon diversity (hypothesis 1) between the control and warmed plots were assessed with generalised linear mixed models using a Bayesian method that relies on Markov Chain Monte Carlo (MCMC) iterations. In these models we treated treatment (control or OTC), *B. nana* abundance and litter abundance as fixed factors and plot as a random factor to account for repeated sampling within plots, using the R package ‘MCMCglmm’ (version 2.29) (Hadfield, 2010). For all models, we used as many iterations as necessary to allow for model convergence and an effective sample size of at least 1000. Interferences of differences between the control and warmed estimates were based on the posterior mode estimates and the 95% Highest Posterior Density Credible Intervals.

We tested the effect of treatment, *B. nana* abundance and litter abundance on the bacterial community composition with PERMANOVAs (Anderson, 2001). All PERMANOVAs were based on Bray-Curtis distance matrices and were performed using the *adonis* function in the R package ‘vegan’ (version 2.5-6). We also tested whether samples taken from the same plot were similar to each other using PERMANOVAs. Plot indeed had a significant effect on the cDNA-based bacterial community composition, but not on the DNA-based bacterial community composition (Table S3 and S4). To reduce possible biases related to samples coming from the same plot, we used plot as *strata* in the PERMANOVAs testing the effect of treatment, *B. nana* abundance and litter abundance. In this way we controlled for the variation caused by repeated sampling within plots by limiting permutations within plots.

The relative abundances of taxa on phylum, class and order level between the warmed and the control samples (hypothesis 2) were tested using Wilcoxon rank-sum tests on plot averages (samples from the same plot were pooled for this purpose) using the *stat_compare_means* function from the R package ‘ggpubr’ (version 0.2.1) (Kassambara, 2020).

Two methods were used to determine taxa on ASV level sensitive to warming (hypothesis 2). First, differential abundance of bacterial genera between warmed and control samples was assessed using the DESeq2 procedure (Love et al., 2014) on the non-CSS normalised datasets (with pseudoreplicates pooled per plot) with the R package ‘DESeq2’ (version 1.24.0) (Love et al., 2014). The adjusted *P*-value cut-off was 0.1 (Love et al., 2014). Differential abundance analysis only uses ASVs present in both the OTC and control samples. The second method we used to find taxa sensitive to warming, was the indicator species analysis. To find bacterial taxa indicative for the warming or the control treatment, correlation-based indicator species analysis was done with all possible site combinations using the function *multipatt* of the R package ‘indicSpecies’ (version 1.7.6) (De Caceres & Legendre, 2009) based on 10^3^ permutations. For this, we pooled all samples originating from the same plot. The indicator species analysis takes into account ASVs present in both OTC and control samples, but also ASVs present in only one of the treatments. We combined results of the DESeq2 and indicator species analysis into a final list of ASVs sensitive to warming. Data are presented as the number of significant ASVs identified in DESeq2 and/or indicator species analysis and represented at the genus level.

To test hypothesis 3, we used structural equation modelling to estimate the direct and indirect effects of warming on the bacterial community and the consequences for N_2_-fixation. The structural equation models were fitted using the R package ‘lavaan’ (version 0.6-7). Initial models were constructed using current knowledge and hypotheses of effects of warming on plant-microbe interactions and on N_2_-fixation activities. As variables included in the model, we used treatment, litter abundance, *B. nana* abundance, 16S rRNA abundance, *nifH* abundance, N_2_-fixation rates and ‘bacterial community structure’. The latter is a latent variable which consisted of the average of β-diversity and Shannon diversity index per plot for the combined cDNA and DNA data. β-diversity was derived from the first axis of a PCoA analysis. All variables were averaged per plot. We tested whether the model has a significant model fit according to the following criteria: χ2/df < 2, P[values (*P* > 0.05), root mean square error of approximation (rmsea) < 0.07 and goodness of fit index (GFI) > 0.9 (Hooper et al., 2008).

## Results

### Treatment effect on bacterial diversity and community structure

The richness and Shannon diversity of the DNA-based and the cDNA-based bacterial communities did not differ significantly between control and OTC samples (Figs 1a-1d, Table S3). However, we found a negative effect of *B. nana* abundance on the richness and Shannon diversity of the cDNA-based bacterial community (richness: pMCMC = 0.004; Shannon diversity pMCMC = 0.01, Figs 1c-1d).

**Figure 1.**
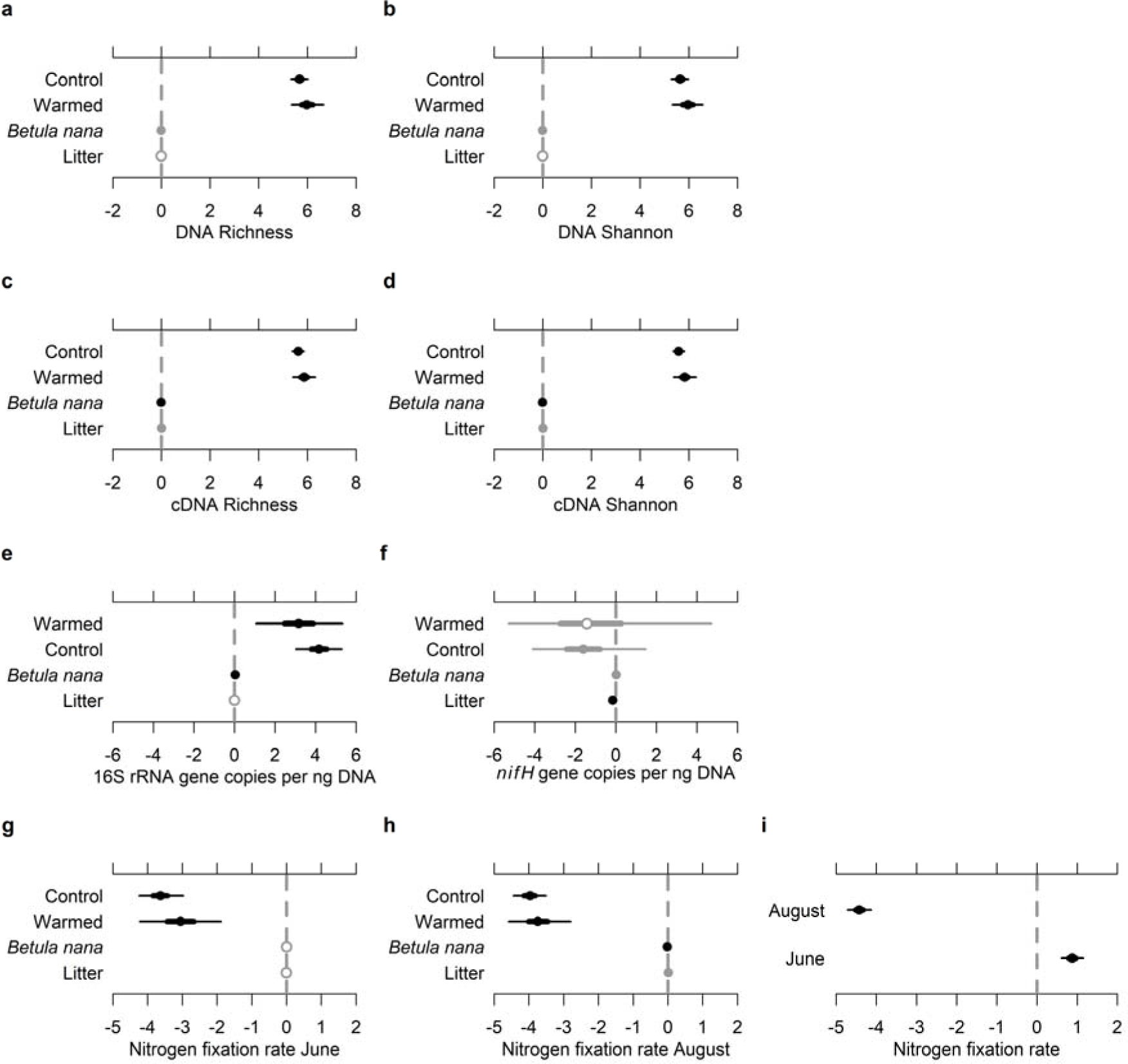
Fixed effect structure of the linear mixed-effect models testing the effect of treatment (warmed and control), *Betula nana* abundance and litter abundance on a) DNA-based richness and b) Shannon diversity, c) cDNA-based richness and d) Shannon diversity, e) 16 rRNA gene abundance, f) *nifH* gene abundance, N_2_-fixation rate g) in June, h) in August and i) fixed effect structure of the linear mixed-effect model testing the difference between N_2_-fixation rates in June and August. Non-overlapping 95% High Posterior Density Credible Interval (95% CrI) are used to detect significant differences between effects. Parameters with 50% CrI overlapping 0 are indicated by open circles. Parameters with 50% CrI not overlapping 0, but with 95% CrI overlapping 0 are indicated by closed black circles. Thick lines represent 50% CrI and thin lines represent 95% CrI.

The PERMANOVA showed that treatment significantly influenced the DNA- and the cDNA-based community compositions of the moss (DNA: R^2^ = 0.05, and *P* < 0.001 and cDNA: R^2^ = 0.04, and *P* < 0.001; Table S6 and S7). In addition to the warming treatment, litter abundance also significantly influenced the DNA-based bacterial community composition (R^2^ = 0.03, *P* = 0.05), but not the cDNA-based bacterial community composition (Table S6 and S7).

### Taxonomic composition R. lanuginosum-associated bacterial communities

In the control samples, where bacterial communities were under ambient environmental conditions, the most abundant phyla in the DNA and cDNA samples included Proteobacteria (44% and 40% average relative abundance across all control DNA and cDNA samples respectively), followed by Acidobacteria (DNA: 29%, cDNA: 23%), Actinobacteria (DNA: 8%, cDNA: 15%), Cyanobacteria (DNA: 7%, cDNA: 2%), Planctomycetes (DNA: 4%, cDNA: 2%), Bacteroidetes (DNA: 4%, cDNA: 4%), Verrucomicrobia (DNA: 2%, cDNA: 3%) and Armatimonadetes (DNA: 2%, cDNA: 2%) (Fig. 2a). The most abundant Proteobacterial class were Alphaproteobacteria (DNA: 29%, cDNA: 31%) (Fig. 2b).

**Figure 2.**
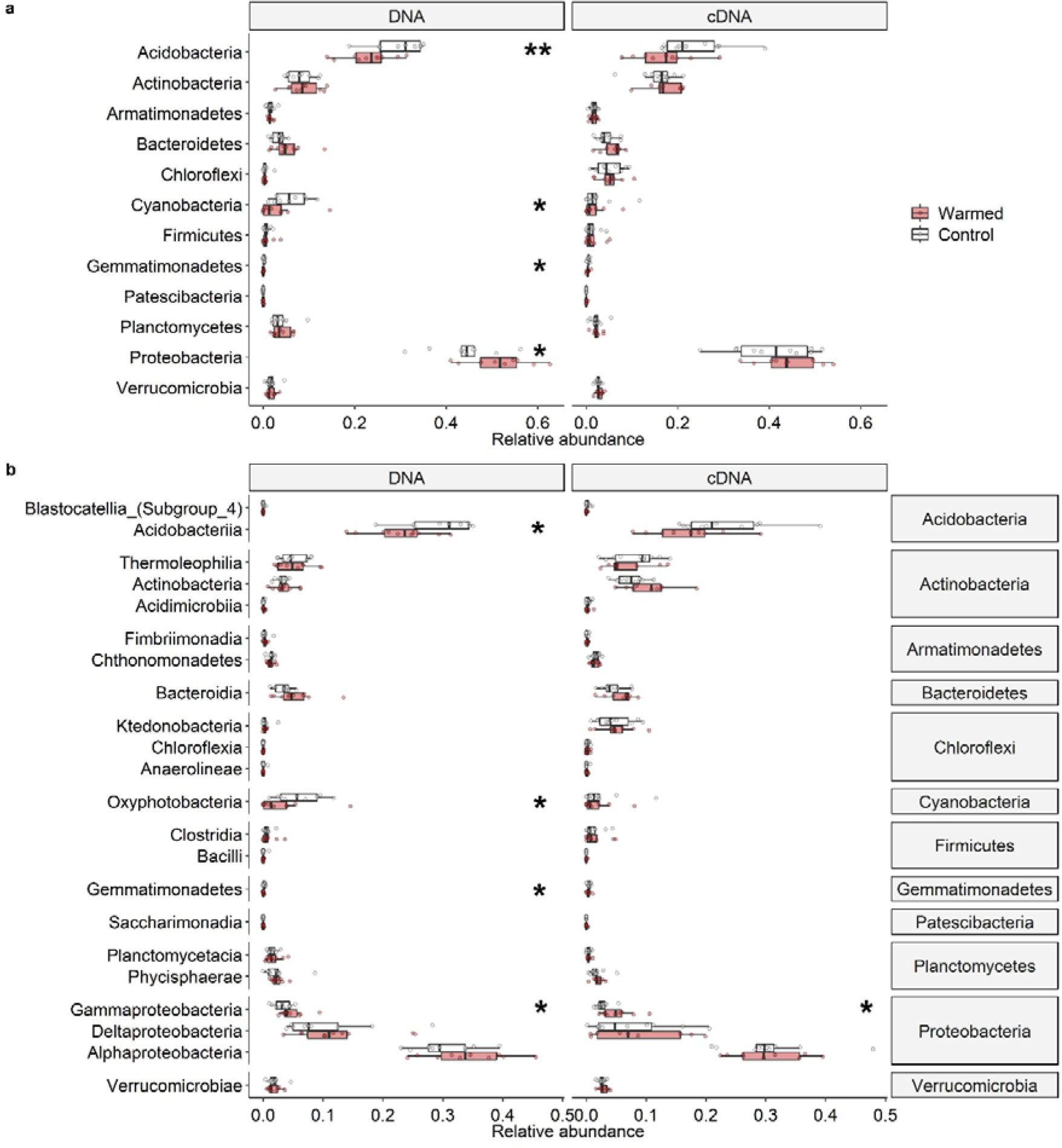
Boxplots of the relative abundances of (A) phyla and (B) classes in DNA and cDNA based bacterial communities associated with the moss *R. lanuginosum*. Boxplots represent minimum values, first quartiles, medians, third quartiles and maximum values. Significance levels (* < 0.05, ** < 0.01) are based on Wilcoxon rank sum tests.

Acetobacterales (DNA: 15%, cDNA: 21%), Myxococcales (DNA: 12%, cDNA: 7%), Caulobacterales (DNA: 6%, cDNA 3%) and Rhizobiales (DNA: 6%, cDNA 5%) were the most abundant orders of the Proteobacteria (Fig. S1). The order Acetobacterales was dominated by the genus *Acidiphilium* (DNA: 5%, cDNA 8%), the order Myxococcales was dominated by the genus *Haliangium* (DNA: 4%, cDNA 3%) (Fig. S2).

The Acidobacteria were dominated by the orders Acidobacteriales (DNA: 17%, cDNA 16%) and Solibacterales (DNA: 11%, cDNA: 7%) (Fig. S1). The Acidobacteriales were dominated by the genus *Granulicella* (DNA: 11%, cDNA: 7%). The Solibacterales were dominated by the genera *Bryobacter* (DNA: 5%, cDNA 2%) and *Candidatus* Solibacter (DNA: 6%, cDNA: 5%) (Fig. S3).

Actinobacteria mainly comprised the orders Solirubrobacterales (DNA: 5%, cDNA: 8%) and Frankiales (DNA: 2%, cDNA: 4%) (Fig. S1).

Cyanobacteria were dominated by the genera *Nostoc* (DNA: 5%, cDNA: 2%) and *Stigonema* (DNA: 1%, cDNA 0.1%) (Fig. S4).

### Treatment effect on the relative abundances of bacterial taxa on phylum, class and order level

We compared the relative abundances of taxa on phylum, class and order level in the controls with the warmed samples from the OTCs (Fig. 2 and Fig. S1). On phylum level, Acidobacteria (Wilcoxon rank-sum test, *P* = 0.008), Cyanobacteria (*P* = 0.03) and Gemmatimonadetes (*P* = 0.02) decreased in relative abundance with warming, while Proteobacteria (*P* = 0.04) increased in relative abundance in the DNA-based bacterial communities (Fig. 2a). We did not detect significant changes in the cDNA-based bacterial communities on phylum level.

On class level, Acidobacteriia (*P* = 0.01), Gemmatimonadetes (*P* = 0.02), and Oxyphotobacteria (*P* = 0.03) decreased in relative abundance under warming in the DNA-based bacterial communities, while Gammaproteobacteria (*P* = 0.04) increased in relative abundance in the DNA- and the cDNA-based bacterial communities (Fig. 2b).

At the order level, Betaproteobacteriales (DNA: *P* = 0.04, cDNA: *P* = 0.005) and Micrococcales (DNA: *P* = 0.007, cDNA: *P* = 0.0007) had a higher relative abundance in the warmed DNA- and cDNA-based bacterial communities (Fig. S1). Acidobacteriales (DNA: *P* = 0.03, cDNA: *P* = 0.04) showed a lower relative abundance in the warmed DNA- and cDNA-based bacterial communities (Fig. S1). In addition, in the DNA-based bacterial communities, Sphingobacterales (*P* = 0.05) and Cytophagales (*P* = 0.02) increased in relative abundance under warming. Nostocales (*P* = 0.03) decreased in relative abundance under warming. In the cDNA-based bacterial communities, the orders Sphingomonadales (*P* = 0.02) and Rhizobiales (*P* = 0.02) increased in relative abundance under warming, while Acetobacterales (*P* = 0.05) decreased in relative abundance under warming (Fig. S1).

### Treatment related shifts in the relative abundance of ASVs

For the bacterial communities in the DNA-based analysis, DESeq2 and indicator species analysis combined revealed 23 ASVs significantly higher in relative abundance under warming and 122 ASVs with higher relative abundance in the controls (Table S8). The strongest indicator species for the control plots corresponded to the taxa that were more abundant in the control plots according the DESeq2 analysis. ASVs with increased relative abundance in the warmed samples belonged to the genera *Allorhizobium*-*Neorhizobium*-*Pararhizobium*-*Rhizobium, Nitrobacter* (Alphaproteobacteria), and *Galbitalea* (Actinobacteria). ASVs with increased relative abundance in the controls belonged to the genera *Acidipila*, *Bryocella*, *Bryobacter*, *Candidatus* Solibacter and *Granulicella* (Acidobacteria), *Acidiphilium*, *Endobacter,* and *Bradyrhizobium* (Alphaproteobacteria), *Nostoc* (Cyanobacteria), and *Conexibacter* (Actinobacteria) (Fig. 3 and Table S8).

**Figure 3.**
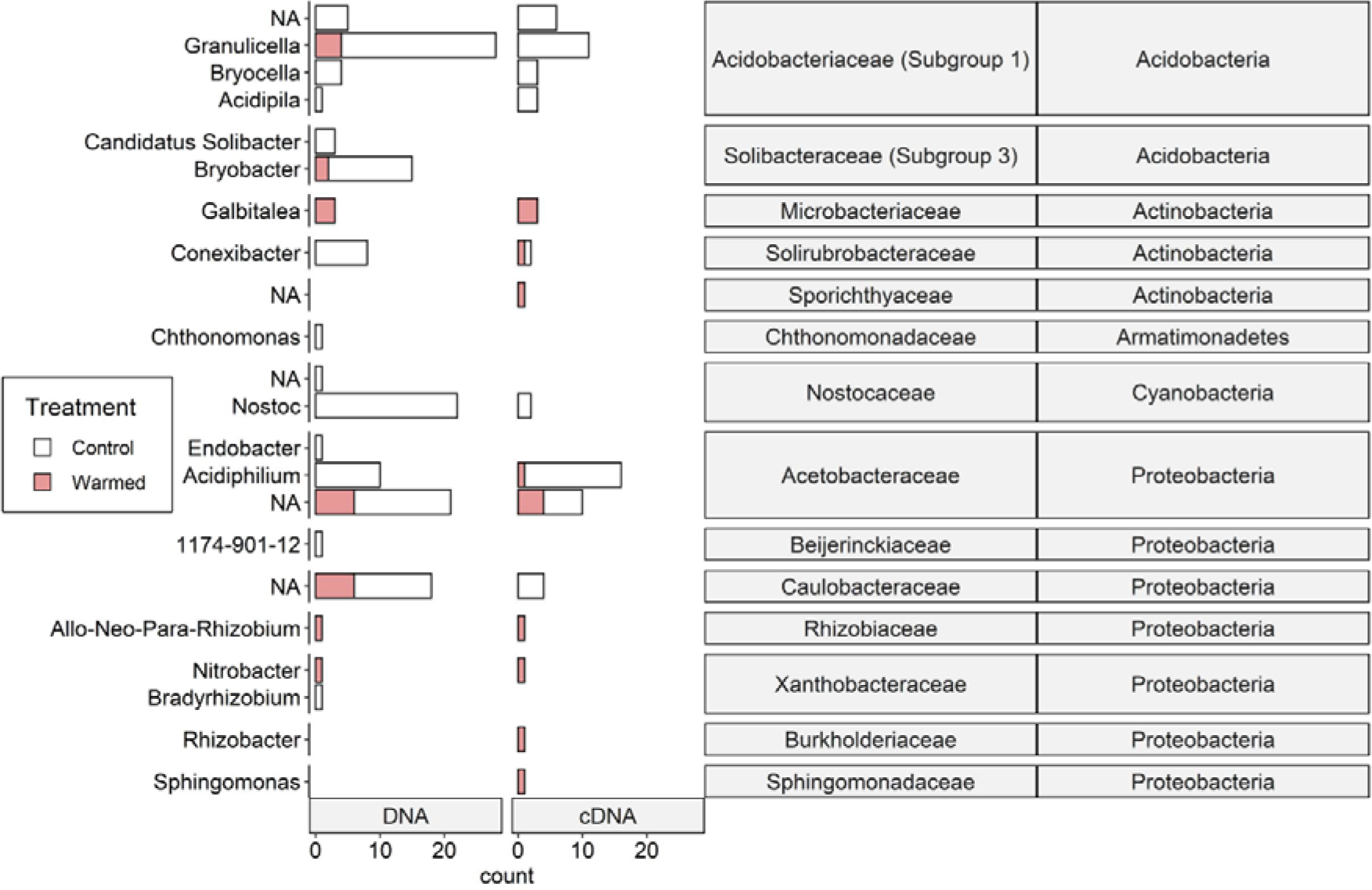
Number of ASVs (amplicon sequence variants) per genus sensitive to warming for DNA and cDNA based bacterial communities associated with the moss *R. lanuginosum*. Sensitivity was determined by differential abundance analysis (DESeq2) and indicator species analysis. ASVs not assigned to genus level are labelled ‘NA’ and ‘Allo-Neo-Para-Rhizobium’ refers to ‘Allorhizobium-Neorhizobium-Pararhizobium-Rhizobium’.

For the bacterial communities in cDNA-based analysis, we detected 54 potentially active ASVs with higher abundance in the control plots and 14 potentially active ASVs more abundant in the warmed plots (Fig. 3, Table S9). ASVs more abundant in the control plots belonged to the genera *Acidipila*, *Bryocella*, *Granulicella* (Acidobacteria), *Nostoc* (Cyanobacteria) and *Acidiphilium* (Alphaproteobacteria). ASVs more abundant under warming belonged to the genera *Allorhizobium*-*Neorhizobium*-*Pararhizobium*-*Rhizobium*, *Nitrobacter*, *Sphingomonas* (Alphaproteobacteria), *Galbitalea* (Actinobacteria), and *Rhizobacter* (Gammaproteobacteria) (Fig. 3, Table S8).

### Treatment effect on 16S rRNA gene and nifH gene copy numbers and nitrogen fixation rates

No significant difference was found in the 16S rRNA gene and *nifH* gene abundance between the control and warmed samples (Figs 1e-1f, Table S3). However, litter abundance negatively affected *nifH* gene abundance (pMCMC = 0.04, Fig. 1f) and *B. nana* abundance tended to positively influence 16S rRNA gene abundance (pMCMC = 0.072, Fig. 1e).

We did not find any differences between N_2_-fixation rates (expressed as produced ethylene) in the control and warmed plots in June or August 2014 (Figs 1g-1h, Table S3). However, N_2_-fixation in control and warmed plots in August was significantly lower (pMCMC < 0.001) than in June (Fig. 1i). We also found a significant negative correlation between *B. nana* abundance and N_2_-fixation activities measured in August (pMCMC = 0.04, Fig. 1h).

### Relationships between treatment, plant biomass, bacterial community structure and N_2_-fixation

To explore the direct and indirect linkages between warming, *B. nana* and litter abundance, bacterial community structure and N_2_-fixation, we constructed a structural equation model (SEM) (Fig. 4, Table S10). We found that warming was directly associated with changes in bacterial community structure and positively correlated with increased of *B. nana* abundance. The direct effect of *B. nana* was stronger than the direct effect of treatment on the bacterial community (-1.2 versus 0.79 standardized regression coefficients). Changes in the bacterial community structure were also indirectly associated with warming through variation in *B. nana* abundance.

**Figure 4.**
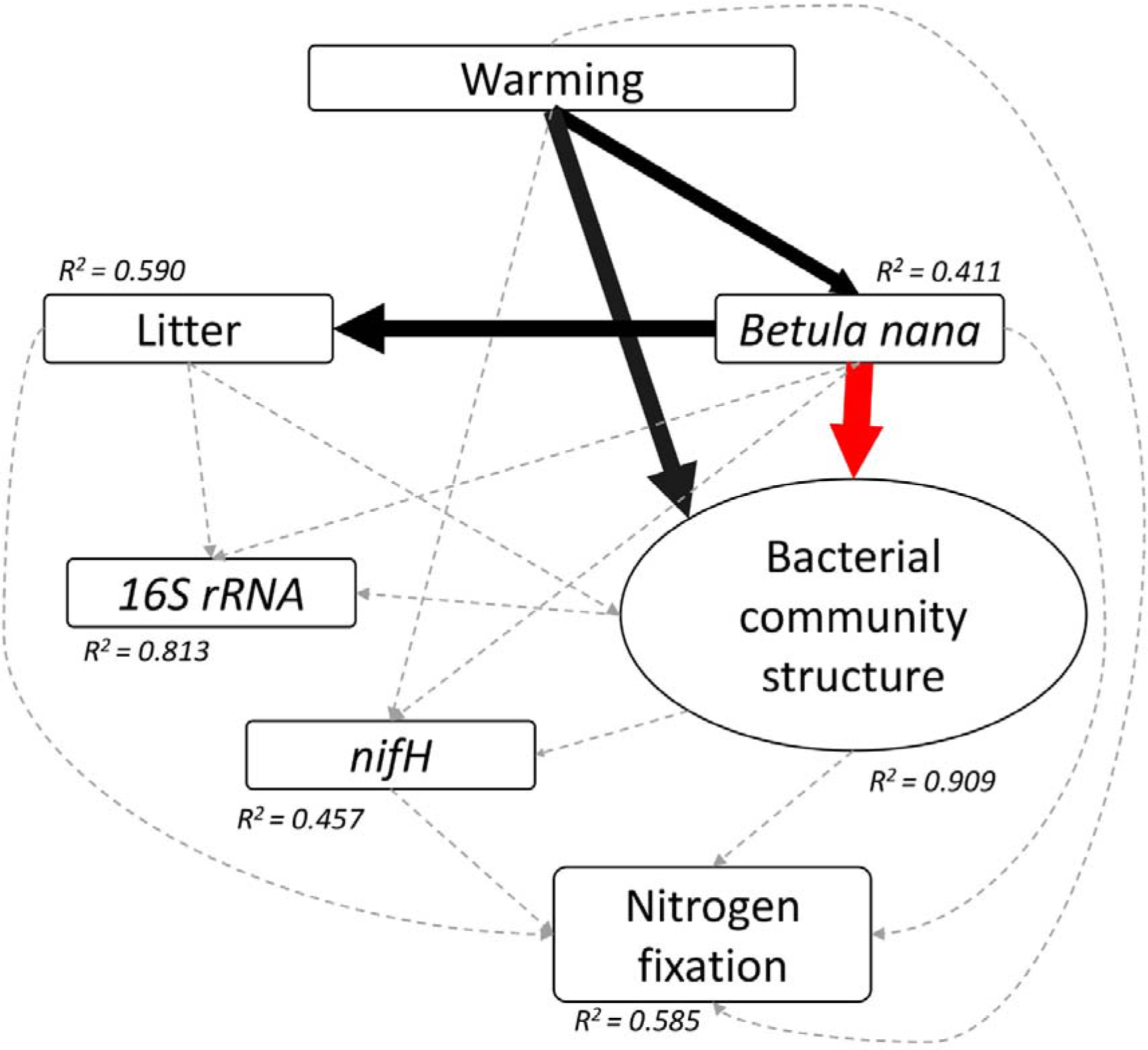
Structural equation model of relationships between warming, *Betula nana* and litter abundance, moss-associated bacterial community and N_2_-fixation. A latent variable (bacterial community structure) was computed to represent β-diversity and Shannon diversity index for each plot. χ2 = 2.896, *P*-value = 0.894, df = 7, GFI = 0.982, RMSEA = 0, TLI = 1.202. Positive significant effects are represented in black and negative significant effects in red. The strength of the effect is visualized by the width of the arrow. The R^2^-value represents the proportion of total variance explained for the specific dependent variable. Dash line arrows indicate non-significant effects. Standardized path coefficients are presented in Table S3.

The strongest positive effect detected was warming treatment on the bacterial community structure and the strongest negative effect was *B. nana* abundance on bacterial community structure (Fig. 4, Table S10).

## Discussion

Mosses form an important C and N sink in high latitudes and their associated bacterial communities are, to a large extent, responsible for N inputs and organic matter decomposition in these environments. Elucidation of the effect of warming on moss-associated bacterial communities will help to understand how climate change affects C and N cycling driven by the bacterial component of mosses in high-latitude ecosystems. We assessed the effect of long-term (20 years) warming by open-top chambers (OTCs) on bacterial communities and N_2_-fixation associated with the moss *R. lanuginosum* at a tundra site in the highlands of Iceland. Overall, our results suggest that moss-associated bacterial communities are sensitive to long-term experimental warming and the associated plant community change, which caused changes in structure and composition. The abundance of bacteria and diazotrophs however, appeared to be unaffected by warming and, consistent with this finding, no effect on N_2_-fixation rates was observed. However, bacterial taxa that benefitted from the warming treatment almost exclusively belonged to groups involved in N-cycling, which might indicate changes in N turnover and usage of this important nutrient for Arctic ecosystem productivity.

### Effect of warming on the moss-associated bacterial community structure

The average temperature increase induced by the OTCs may seem small (1-2°C), but a temperature increase in this range can affect microbial growth rate, respiration, C uptake and turnover (Walker et al., 2018). In addition, the effect of the OTC treatment is a long-term (20-year) disturbance, which has shown a clear effect on the vegetation structure and biomass (Jonsdottir et al., 2005) and thereby also leads to indirect effects of warming on the microbial community.

The richness and Shannon diversity of the total and potentially metabolically active bacterial community were not significantly affected by 20 years of warming. These results contrast with our first hypothesis and with trends of decreasing richness and diversity in *Sphagnum* moss observed by Carrell et al. (2017) and by Kolton et al. (2019). *R. lanuginosum* has a much lower water holding capacity than *Sphagnum* (Elumeeva et al., 2011), a different physiology and grows in heathlands and therefore *R. lanuginosum* might react differently to warming. In addition, while our study describes the effect of 20 years warming *in situ*, those previous studies on *Sphagnum* were much shorter such as a four week laboratory (Kolton et al., 2019) and two years *in situ* experimental warming study (Carrell et al., 2019). Nevertheless, we found that warming altered the bacterial community structure, even though only a small part of the variation could be directly explained by the warming treatment. Warming correlated with an increase in shrub and litter abundance and a decrease in moss abundance, as already observed in the site after 3-4 years of warming (Jonsdottir et al., 2005). Indeed, a small part of the variation of the total bacterial community could be attributed to litter abundance, which also negatively affected the richness and diversity of the potentially active bacterial community. In addition, the SEM showed that the bacterial community structure was indirectly correlated with warming via changes in *B. nana* abundance, and indirectly via the combined effect of *B. nana* and litter abundance. The effect of the increase in *B. nana* abundance as result of warming was stronger than the direct effect of warming on the bacterial community structure.

Warming-induced changes in environmental factors such as lower moss layer thickness, higher soil organic matter content, lower soil moisture (Björnsdóttir, 2018; Jonsdottir et al., 2005), or other not measured variables such as leaf nutrient content (Koyama et al., 2018; Sayer et al., 2017; Vandenkoornhuyse et al., 2015) could also contribute to the variation in bacterial communities between moss shoots.

We did not find an effect of warming on the 16S rRNA gene abundance, but *B. nana* abundance was correlated with an increase in 16S rRNA gene abundance. However, as we are not sure about the degree of bias towards chloroplast and mitochondrial DNA of the 16S rRNA gene primers in our samples, we cannot conclude that the bacterial load is indeed affected by *B. nana* abundance.

### Effect of warming on moss-associated bacterial taxa

The total and potentially active bacterial community of *R. lanuginosum* was dominated by Proteobacteria and Acidobacteria, whereas Actinobacteria, Cyanobacteria, Planctomycetes, Bacteroidetes and Verrucomicrobia were present in lower abundances. In agreement with the bacterial community composition of boreal moss species (Holland Moritz et al., 2018) and *Sphagnum* species (Bragina, Berg, et al., 2012), *R. lanuginosum* also showed a high abundance of the Proteobacterial order Acetobacerales and the Acidobacterial order Acidobacteriales.

We analysed changes in relative abundances in several ways to better understand the warming response of the moss bacterial community. This revealed changes in the relative abundances of taxa on phylum, class, order and ASV levels. We hypothesized that the warming-induced increase in labile *B. nana* litter (Jonsdottir et al., 2005) would lead to a decrease in slow-growing, more oligotrophic taxa, while fast-growing copiotrophic taxa would increase in relative abundance. Our data show indications for a decrease in the relative abundance of oligotrophic taxa in response to warming, such as Acidobacteria (and more specific ASVs of the genera *Granulicella*, *Solibacter*, *Bryocella*, *Bryobacter* and *Acidipila*) (Dedysh & Sinninghe Damsté, 2018; Fierer et al., 2007) and the Alphaproteobacterial genus *Acidiphilium* (Hiraishi & Imhoff, 2015). Acidobacteria often dominate tundra soils (Männistö et al., 2013), especially environments with high concentrations of phenolic compounds, (for instance in *Sphagnum* peat (Pankratov et al., 2011) and *Empetrum* heath (Gallet et al., 1999; Männistö et al., 2013)). In shrub tundra dominated by *B. nana* and *Salix* species, Proteobacteria dominate the soil bacterial community (Wallenstein et al., 2007). In our study, the increase in the relative abundance of Proteobacteria (more specifically the genera *Rhizobacter*, *Nitrobacter* and *Rhizobium*) associated with *R. lanuginosum* in the warmed plots could thus be due to the increase in dwarf shrub biomass and labile litter, selecting for copiotrophic taxa, such as Rhizobiales (Starke et al., 2016). Some oligotrophic taxa with increased abundance in the warmed conditions such as Sphingomonadales and ASVs of the Caulobacterales (Garrity et al., 2015) could be involved in degradation of more recalcitrant plant organic matter (McGenity, 2019; Starke et al., 2016). Caulobacterales has for instance been shown to be able to degrade lignin (Wilhelm et al., 2019), which can be found in high concentrations in *B. nana* roots and leaves (McLaren et al., 2017). An increase in *B. nana* litter likely increases the rate of C fluxes (Parker et al., 2018), and this may partly be due to a shift towards faster growing copiotrophic bacterial taxa, at least in the moss layer.

While the overall warming-induced changes in bacterial phylotypes for the total and the potentially active bacterial community were similar, we found that the total bacterial community reacted more strongly to warming than the potentially active bacterial community in terms of changes in relative abundance of the number of phyla, classes and ASVs. This difference may be explained by a difference in drivers for the total and potentially metabolically active bacteria, with changes in total bacterial community structure reflecting long-term drivers, while the active bacterial community may reflect short-term differences between OTC and controls (Y. Wang et al., 2020).

### Implications of warming for the moss bacterial community involved in N-cycling

Our results of the bacterial structure and composition revealed that warming induced changes in relative abundances of several taxa potentially involved in N-cycling. Here, it appears that these taxa involved in the first steps of the N-cycle (entrance of new N through N_2_-fixation and production of nitrate from nitrite) are altered by warming. Although we did not explicitly target the N_2_-fixing or nitrifying community by sequencing in this study, we found indications for changes in the relative abundance of potentially N_2_-fixing and nitrifying taxa. In particular, the relative abundance of Cyanobacteria decreased. At the genus level, this was characterized by the lower abundance of the genus *Nostoc*. The vast majority of taxa that exclusively increased in abundance and had a higher potential metabolic activity under warming belong to groups capable of N_2_-fixation (*Sphingomonas*, *Allorhizobium*-*Neorhizobium*-*Pararhizobium*-*Rhizobium*, *Rhizobacter*) and nitrification (*Nitrobacter*). However, neither *nifH* gene abundance nor N_2_-fixation rates were directly affected by warming. This is in agreement with the general response of N_2_-fixation and abundances of *nifH* genes to warming in cold ecosystems (Salazar et al., 2019), where N_2_-fixation and abundances of *nifH* genes are unresponsive and nitrification rates increase under warming treatments. The apparent lack of response in N_2_-fixation rates to warming may be due to a combination of several direct and indirect effects of warming on N_2_-fixation rates counterbalancing each other: the shift in the potentially N_2_-fixing community towards taxa better adapted to the new environment and thereby compensating for the decrease in Cyanobacteria in our study, the negative effect of drier conditions due to the warming treatment (Rousk et al., 2018; Whiteley & Gonzalez, 2016), the direct positive effect of warming (Rousk & Michelsen, 2017) and the negative effect of shading due to increasing shrub cover (Sorensen et al., 2012) and the positive or negative effect of fertilization by shrub litter (Rousk & Michelsen, 2017; Sorensen & Michelsen, 2011). The SEM however did not indicate any links between the warming treatment, the bacterial community structure, litter and *B. nana* abundance to *nifH* gene abundance or N_2_-fixation rates. One reason for this could be that a degree of functional compensation occurs through the shift in the diazotrophic community with warming. Nevertheless, the findings are supporting hypothesis 3, and *nifH* gene abundance was negatively affected by litter abundance and N_2_-fixation rates in August were negatively affected by *B. nana* abundance, indicating the presence of indirect effects of warming on N_2_-fixation. N_2_-fixation rates in August were also lower than in June in both OTCs and control plots, maybe due to an increase of the effect of shading in August as indicated by the effect of *B. nana*. It may also have been drier in August, or it could be due to a seasonal shift in the N_2_-fixing bacterial community (Warshan et al., 2016).

Finally, it is important to note that *R. lanuginosum* biomass tends to decrease in the warmed plots (Björnsdóttir, 2018; Jonsdottir et al., 2005, Table S2). Thus considering that in our study N_2_-fixation rates are expressed per gram moss, warming would consequently lead to a reduction of the total amount of N_2_ fixed per unit area in this tundra ecosystem.

Our study is among the first to assess the effect of long-term (20 years) experimental warming with OTCs on the bacterial part of a moss microbiome. Our results showed no direct response of N_2_-fixation rates and *nifH* gene abundance to warming. However, long-term warming led to changes in the bacterial community composition. On ASV level, these changes were characterized by a decrease in the relative abundance of Cyanobacteria and an increase in abundance and potential metabolic activity of non-cyanobacterial diazotrophs, which may explain the lack of response of N_2_-fixation to warming. Our results also showed that warming-induced changes in the surrounding vegetation structure can affect moss-associated bacterial communities, thus underlining the importance of indirect effects of long-term warming. The bacterial community associated with the moss might thus be sensitive to future warming, with potential implications for N_2_-fixation rates, moss growth and C sequestration.

## Funding

This work was supported by the MicroArctic Innovative Training Network grant supported by the European Commissions’s Horizon 2020 Marie Sklodowska-Curie Actions program [grant number 675546] and the Energy Research Fund [grants to ISJ, numbers 08-2013, NÝR-09-2014].

## Acknowledgements

We thank Dr. Ólafur S. Andrésson, Dr. Marie-Charlotte Nilsson, and Dr. Matthias Zielke for help and advice on the RNA and DNA extractions and the ARAs. We would also like to thank Quentin J.B. Horta-Lacueva for advice on the statistical analysis.

## Competing Interests

The authors declare that the research was conducted in the absence of any conflict of interest.

## Author contributions

IJK, AJRC, ISJ and OV designed the study. IJK, AJRC and CK performed the research. IJK analysed the data and wrote the paper with input from AJRC, CK, DW, ADJ, ISJ and OV.

## Data availability

Raw sequences are available in the European Nucleotide Archive under accession number PRJEB40635.

## Notes

### Competing Interest Statement

The authors have declared no competing interest.

