## Supplementary Material for "Long-term warming effects on the microbiome and nitrogen fixation of a common moss species in sub-Arctic tundra"

**Figure S1.** Boxplots of the relative abundances of bacteria orders associated with the moss *R. lanuginosum* at the order level for DNA- and cDNA-based bacterial community samples associated with the moss *R. lanuginosum*. Controls are shown in white and OTC (warmed) samples are shown in red. Boxplots represent minimum values, first quartiles, medians, third quartiles and maximum values. Significance levels ( * < 0.05, ** < 0.01, *** < 0.001) are based on Wilcoxon rank sum tests.

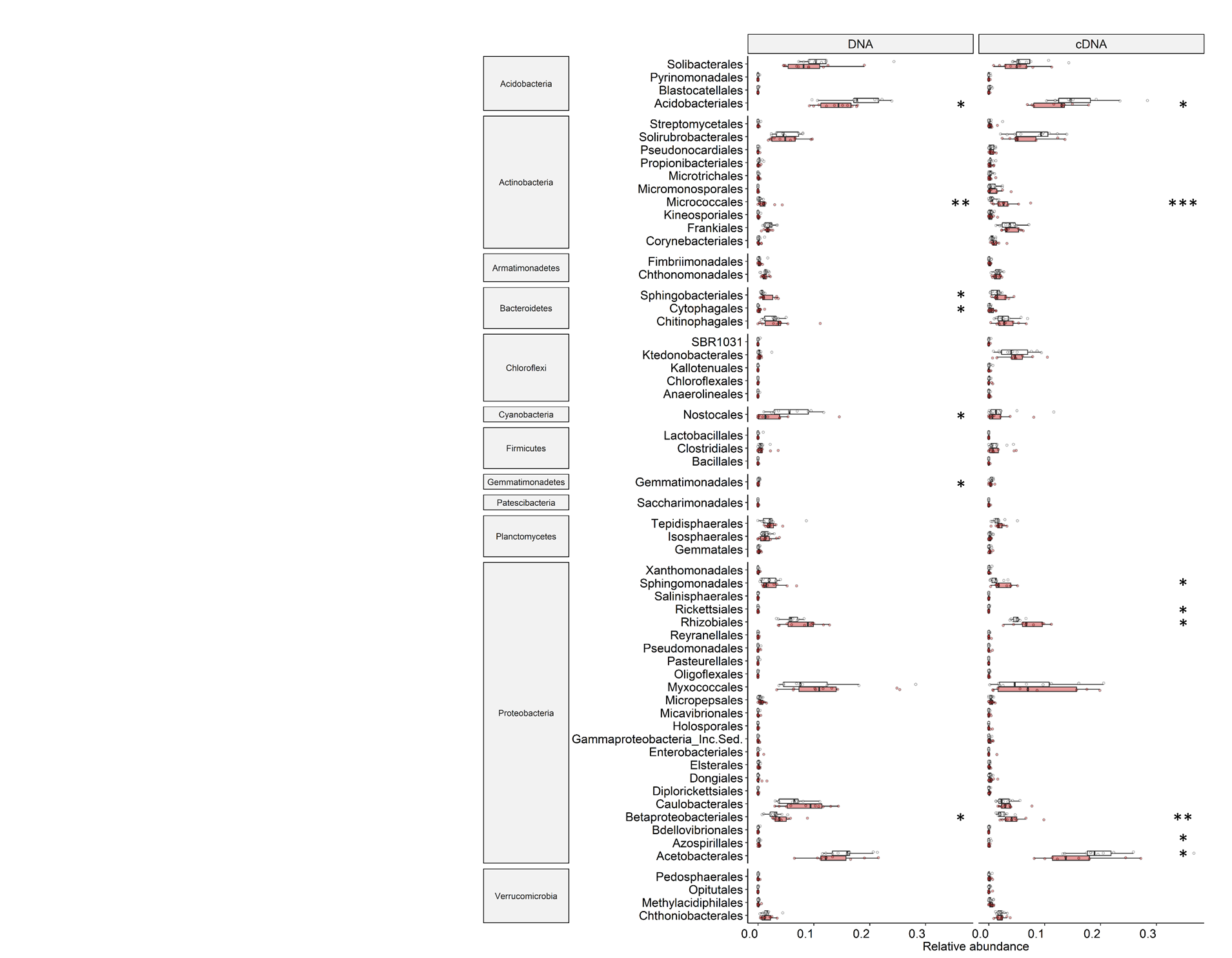

**Figure S2** Barplots showing the relative abundance of Alphaproteobacterial genera in warmed and control plots of the cDNA- and the DNA-based bacterial communities associated with the moss *R. lanuginosum*.
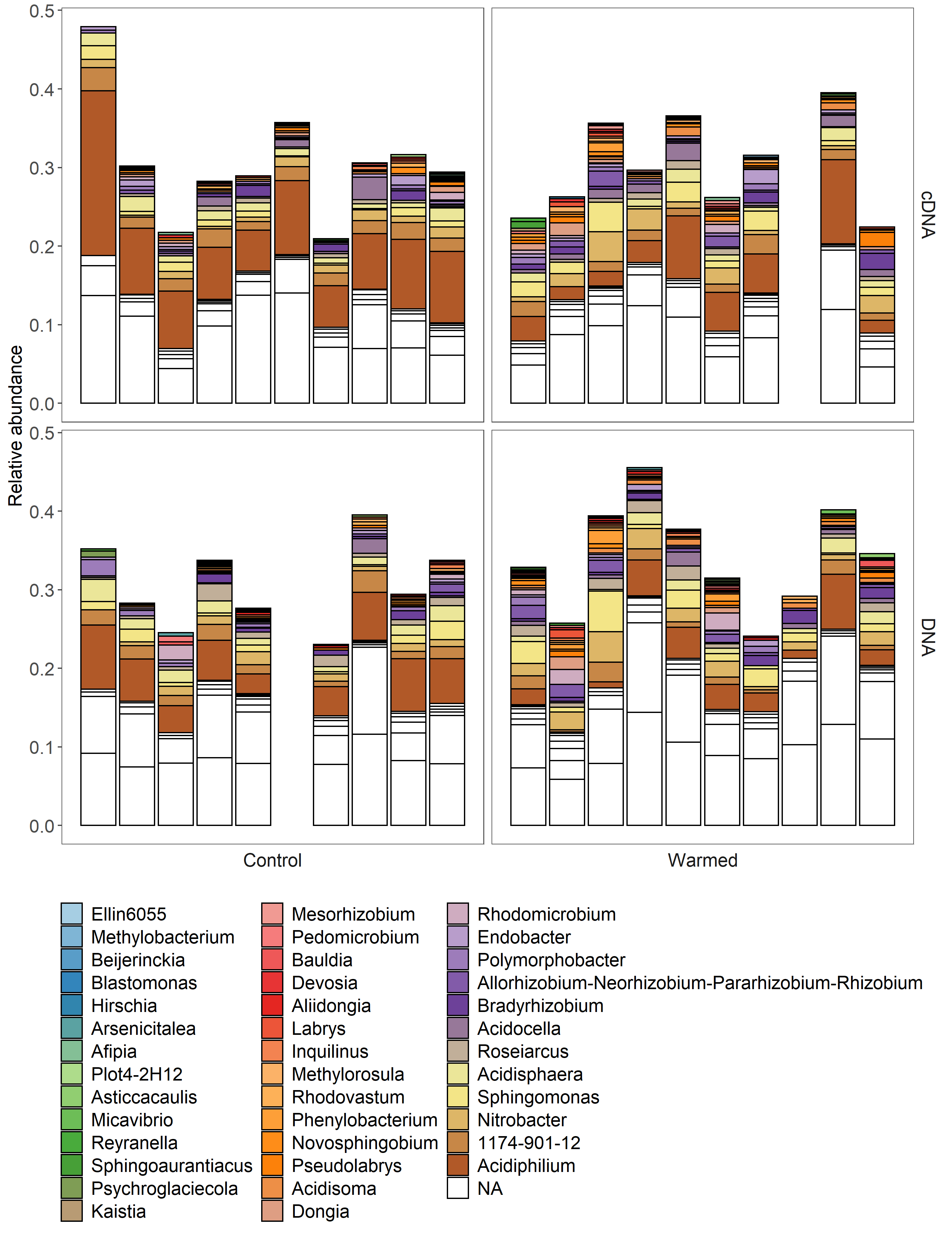

**Figure S3** Barplots showing the relative abundance of Acidobacterial genera in warmed and control plots of the cDNA- and the DNA-based bacterial communities associated with the moss *R. lanuginosum*.

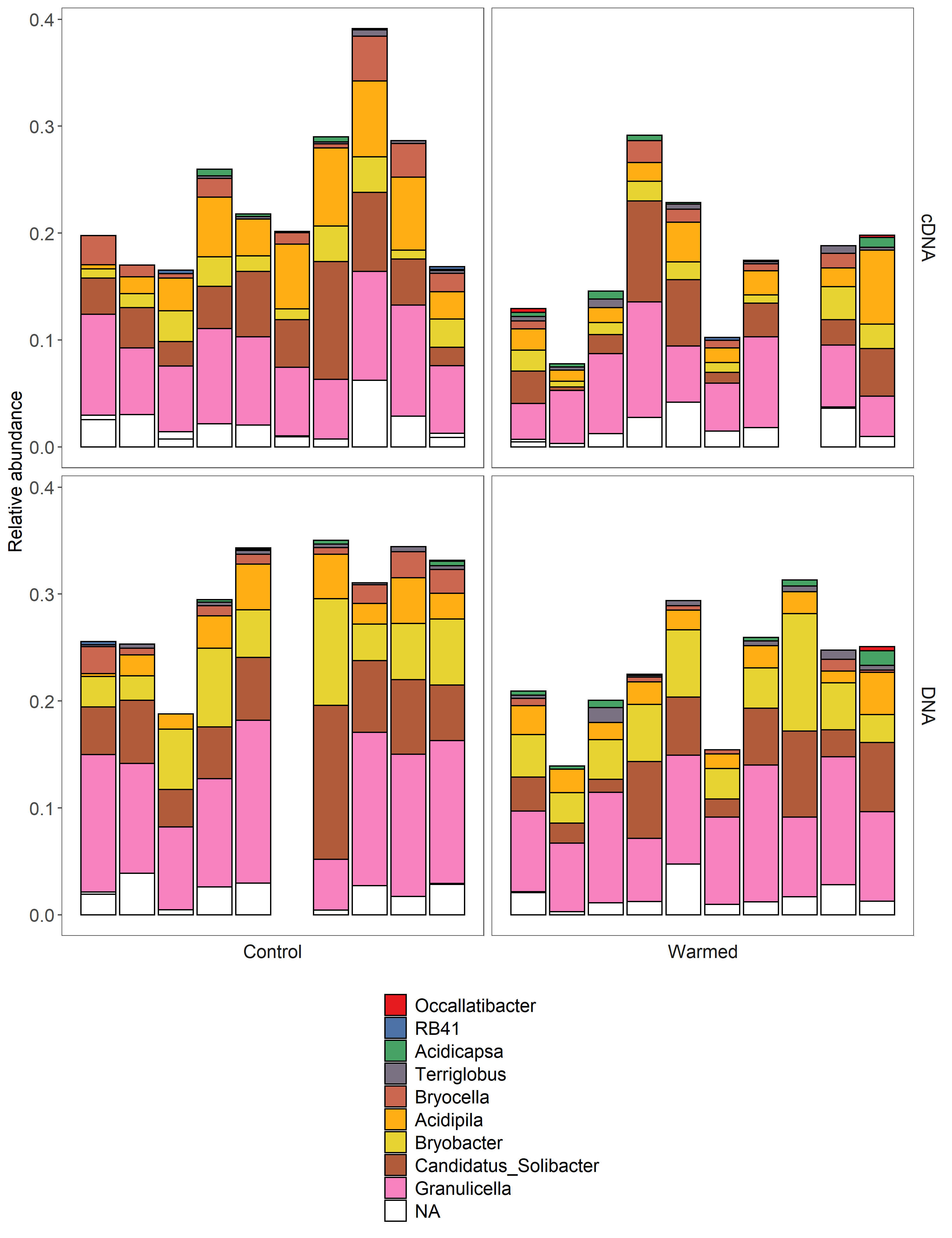

**Figure S4** Barplots showing the relative abundance of Cyanobacterial genera in warmed and control plots of the cDNA- and the DNA-based bacterial communities associated with the moss *R. lanuginosum*.

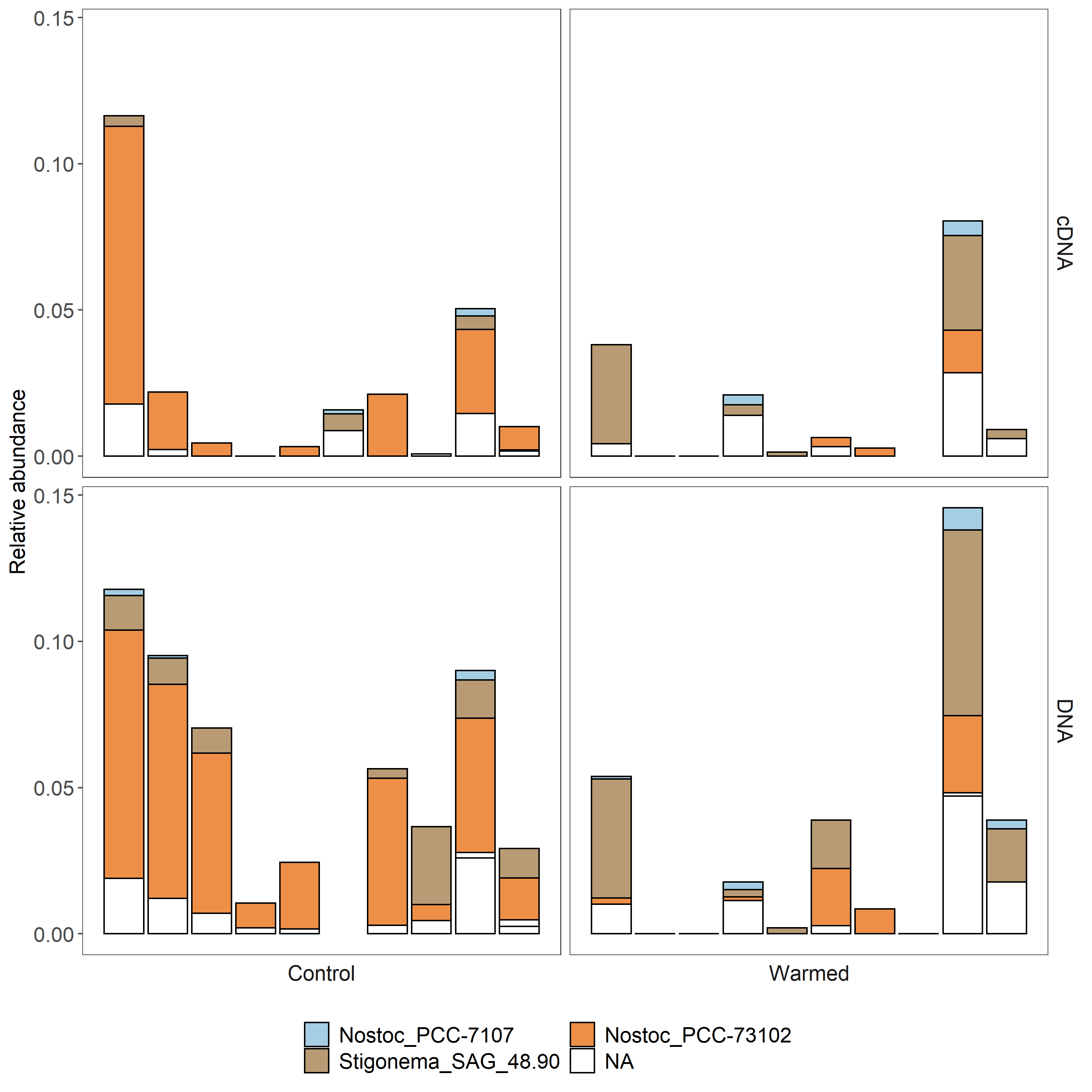

**Table S1.** Temperature and relative humidity for the OTC (warmed) and control plots measured in June-August 2016 (temperature and relative humidity 10 cm above the moss layer) and August 2018-June 2019 (temperature on the moss surface). Shown are mean ± standard error of the mean. Significant differences (t-test, *P* < 0.05) are indicated in bold.

| Air temperature  June – August 2016 | | | Moss surface temperature  August 2018 – June 2019 | | | Relative humidity Air  June – August 2016 | | |
| --- | --- | --- | --- | --- | --- | --- | --- | --- |
| OTC | Control | ∆°C | OTC | Control | ∆°C | OTC | Control | ∆% |
| 11.4  ± 0.1 | 10.0  ± 0.1 | **1.4** | 1.28  ± 0.01 | 1.06  ± 0.01 | **0.22** | 78.8  ± 0.36 | 81.8  ± 0.37 | **-3** |

**Table S2.** Abundance (total hits) for *Racomitrium lanuginosum*, *Betula nana* and litter in controls and OTCs. Differences between controls and OTCs were tested with paired t-tests. ** p<0.01, *** p<0.001. ). Shown are mean ± standard error of the mean. Significant differences (t-test, *P* < 0.05) are indicated in bold.

| *Racomitrium lanuginosum* | | | *Betula nana* | | | Litter | | |
| --- | --- | --- | --- | --- | --- | --- | --- | --- |
| OTC | Control | ∆ | OTC | Control | ∆ | OTC | Control | ∆ |
| 48.1  ± 6.53 | 58.7  ± 4.68 | -10.6 | 59.7  ± 7.09 | 23.8  ± 5.54 | **35.9** | 23.1  ± 4.00 | 8.7  ± 1.96 | **14.4** |

| **Table S3.** Bacterial richness and diversity indicators, 16S rRNA and *nifH* gene copy numbers per ng DNA, and N_2_-fixation rates in June and August in control and warmed plots (OTC). Shown are mean ± standard error of the mean. | | |
| --- | --- | --- |
|  | Control | OTC |
| DNA richness | 267.4 ± 36.6 | 240.2 ± 29.3 |
| DNA Shannon diversity | 5.38 ± 0.13 | 5.29 ± 0.12 |
| cDNA richness | 248.3 ± 25.7 | 206.1 ± 13.0 |
| cDNA Shannon diversity | 5.37 ± 0.10 | 5.26 ± 0.06 |
| 16S rRNA gene copy number per ng DNA | 2681 ± 1313 | 3902 ± 1463 |
| *nifH* gene copy number per ng DNA | 17.4 ± 8.3 | 4.5 ± 2.1 |
| N_2_-fixation rate June | 0.040 ± 0.0092 | 0.039 ± 0.0035 |
| N_2_-fixation rate August | 0.018 ± 0.0022 | 0.014 ± 0.0019 |

| **Table S4.** Summary table for the Permanova testing the effect of plot on the DNA-based bacterial community variation of the moss. | | | | | | |
| --- | --- | --- | --- | --- | --- | --- |
| Source | Df | Sum of Squares | Mean Squares | F | R^2^ | P |
| Plot | 1 | 0.4982 | 0.49823 | 1.2182 | 0.02886 | 0.09191 |
| Residuals | 41 | 16.7681 | 0.40898 |  | 0.97114 |  |
| Total | 42 | 17.2663 |  |  | 1 |  |

| **Table S5.** Summary table for the Permanova testing the effect of plot on the DNA-based bacterial community variation of the moss. | | | | | | |
| --- | --- | --- | --- | --- | --- | --- |
| Source | Df | Sum of Squares | Mean Squares | F | R^2^ | P |
| Plot | 1 | 0.5932 | 0.59320 | 1.4692 | 0.03543 | 0.01199 |
| Residuals | 40 | 16.1502 | 0.40375 |  | 0.96457 |  |
| Total | 41 | 16.7434 |  |  | 1 |  |

| **Table S6.** Summary table for the Permanova testing the effect of treatment, *Betula nana* abundance and litter abundance on the DNA-based bacterial community variation of the moss. | | | | | | |
| --- | --- | --- | --- | --- | --- | --- |
| Source | Df | Sum of Squares | Mean Squares | F | R^2^ | P |
| Treatment | 1 | 0.7719 | 0.77194 | 1.9574 | 0.04471 | 1.00E-04 |
| *Betula nana* | 1 | 0.5489 | 0.54891 | 1.3919 | 0.03179 | 0.2978 |
| Litter | 1 | 0.5654 | 0.56544 | 1.4338 | 0.03275 | 0.0448 |
| Residuals | 39 | 15.38 | 0.39436 |  | 0.89075 |  |
| Total | 42 | 17.2663 |  |  | 1 |  |

| **Table S7.** Summary table for the Permanova testing the effect treatment, *Betula nana* abundance and litter abundance on the cDNA-based bacterial community variation of the moss. | | | | | | |
| --- | --- | --- | --- | --- | --- | --- |
| Source | Df | Sum of Squares | Mean Squares | F | R^2^ | P |
| Treatment | 1 | 0.694 | 0.69401 | 1.7769 | 0.04145 | 1.00E-04 |
| *Betula nana* | 1 | 0.7071 | 0.70706 | 1.8103 | 0.04223 | 0.7089 |
| Litter | 1 | 0.5 | 0.50002 | 1.2802 | 0.02986 | 0.2587 |
| Residuals | 38 | 14.8423 | 0.39059 |  | 0.88646 |  |
| Total | 41 | 16.7434 |  |  | 1 |  |

| Table S10. Summary statistics of the structural equation model of direct and indirect effects of warming on N_2_-fixation shown in Figure 6. Shown are the standardized path coefficients (Std. est.), the standard error of regression weight (se), the z-value (z) and the significance level for the regression weight (p). | | | | |
| --- | --- | --- | --- | --- |
| Parameter | Std. est. | se | z | p |
| *Microbial community*  *(Shannon diversity and position on first PCoA axis)* |  |  |  |  |
| Treatment | **0.79** | **0.249** | **3.176** | **0.001** |
| B. nana | **-1.208** | **0.314** | **-3.85** | **0** |
| Litter | -0.043 | 0.345 | -0.124 | 0.901 |
| *B. nana* |  |  |  |  |
| Treatment | **0.641** | **0.09** | **7.11** | **0** |
| *Litter* |  |  |  |  |
| B. nana | **0.768** | **0.069** | **11.21** | **0** |
| *16S rRNA gene abundance* |  |  |  |  |
| B. nana | -1.967 | 3.21 | -0.613 | 0.54 |
| Litter | 0.441 | 0.671 | 0.658 | 0.511 |
| Treatment | 1.067 | 2.11 | 0.506 | 0.613 |
| Microbial community | -1.939 | 2.463 | -0.787 | 0.431 |
| *nifH gene abundance* |  |  |  |  |
| B. nana | -1.749 | 1.919 | -0.911 | 0.362 |
| Litter | -0.044 | 0.571 | -0.077 | 0.939 |
| Treatment | 0.677 | 1.272 | 0.532 | 0.595 |
| Microbial community | -1.586 | 1.433 | -1.107 | 0.268 |
| *N_2_-fixation rate (June)* |  |  |  |  |
| Treatment | 1.572 | 4.052 | 0.388 | 0.698 |
| Microbial community | -2.667 | 5.332 | -0.5 | 0.617 |
| B. nana | -3.194 | 6.602 | -0.484 | 0.629 |
| Litter | -0.241 | 0.937 | -0.257 | 0.797 |
| *nifH* gene abundance | -0.702 | 0.816 | -0.861 | 0.39 |
| *Indirect effects on the microbial community and on N_2_-fixation* |  |  |  |  |
| Treatment → Microbial community +  Treatment → B. nana → Microbial community +  Treatment → B. nana → litter → Microbial community | -0.006 | 0.275 | -0.021 | 0.983 |
| Treatment → B. nana → Microbial community +  Treatment → B. nana → litter → Microbial community | **-0.795** | **0.198** | **-4.019** | **0** |
| Treatment → B. nana → Microbial community | **-0.774** | **0.26** | **-2.975** | **0.003** |
| Treatment → B. nana → Litter → Microbial community | -0.021 | 0.17 | -0.124 | 0.901 |
| Treatment → N_2_-fixation +  Treatment → Microbial community → N_2_-fixation +  Treatment → B. nana → Microbial community → N_2_-fixation +  Treatment → B. nana → N_2_-fixation +  Treatment → B. nana → Litter → N_2_-fixation +  Treatment → B. nana → Litter → Microbial community → N_2_-fixation +  Treatment → nifH gene abundance → N_2_-fixation +  Treatment → B. nana → Litter → Microbial community → nifH gene abundance → N_2_-fixation +  Treatment → B. nana → Litter → nifH gene abundance → N_2_-fixation +  Treatment → B. nana → nifH gene abundance → N_2_-fixation | -0.275 | 0.391 | -0.702 | 0.483 |
| Treatment → Microbial community → N_2_-fixation +  Treatment → B. nana → Microbial community → N2-fixation +  Treatment → B. nana → N_2_-fixation +  Treatment → B. nana → Litter → N_2_-fixation +  Treatment → B. nana → Litter → Microbial community → N_2_-fixation +  Treatment → nifH gene abundance → N_2_-fixation +  Treatment → B. nana → Litter → Microbial community → nifH gene abundance → N_2_-fixation +  Treatment → B. nana → Litter → nifH gene abundance → N_2_-fixation +  Treatment → B. nana → nifH gene abundance → N_2_-fixation | -1.846 | 4.122 | -0.448 | 0.654 |
| Treatment → Microbial community → N_2_-fixation | -2.106 | 4.39 | -0.48 | 0.631 |
| Treatment → B. nana → Microbial community → N_2_-fixation | 2.065 | 4.312 | 0.479 | 0.632 |
| Treatment → B. nana → N_2_-fixation | -2.046 | 4.244 | -0.482 | 0.63 |
| Treatment → B. nana → Litter → N_2_-fixation | -0.119 | 0.462 | -0.257 | 0.797 |
| Treatment → B. nana → Litter → Microbial community → N_2_-fixation | 0.056 | 0.468 | 0.12 | 0.904 |
| Treatment → nifH gene abundance → N_2_-fixation | -0.475 | 1.283 | -0.37 | 0.711 |
| Treatment → B. nana → Litter → Microbial community → nifH gene abundance → N_2_-fixation | -0.023 | 0.194 | -0.121 | 0.904 |
| Treatment → B. nana → Litter → nifH gene abundance → N2-fixation | 0.015 | 0.199 | 0.076 | 0.939 |
| Treatment → B. nana → nifH gene abundance → N_2_-fixation | 0.787 | 1.576 | 0.499 | 0.618 |
